# Behavioral and Eye-Tracking Evidence for Disrupted Event Segmentation during Continuous Memory Encoding Due to Short Video Watching

**DOI:** 10.1101/2024.08.17.608429

**Authors:** Hongxiao Li, Jiashen Li, Xin Hao, Wei Liu

**Affiliations:** Key Laboratory of Adolescent Cyberpsychology and Behavior (CCNU), Ministry of Education, Wuhan, China; Key Laboratory of Human Development and Mental Health of Hubei Province, School of Psychology, Central China Normal University, Wuhan, China

**Author notes:** Correspondence: Dr. Xin Hao, School of Psychology, Central China Normal University Address: the 8^th^ floor, Nanhu Complex Building, No 152 Luoyu Road, Wuhan. Postal Code: 430079, Dr. Wei Liu, School of Psychology, Central China Normal University Address: the 8^th^ floor, Nanhu Complex Building, No 152 Luoyu Road, Wuhan. Postal Code: 430079.

**Keywords:** Short Video, Event Segmentation, Eye-tracking, Intersubject Correlation, Hidden Markov Model

## Abstract

The proliferation of short-video platforms prompts critical investigation of their effects on human cognitive functions. We hypothesized that the frequent, user-driven content shifts inherent to short-video watching impair event segmentation—a cognitive process critical for organizing continuous experience into discrete, episodic memory. To investigate this hypothesis, we combined behavioral memory tasks, eye-tracking, and self-report questionnaires. Study 1 (N=113) revealed that exposure to randomly selected short videos impaired subsequent memory for continuous movies. This impairment was not observed following exposure to personalized short videos, nor was it present in trial-based static image encoding tasks (Study2, N=60), suggesting a selective disruption of continuous memory encoding. Intersubject correlation (ISC) analysis of eye movements revealed decreased synchronization at event boundaries during movie watching after exposure to random short videos. Furthermore, the Hidden Markov Model (HMM) analysis indicated that this exposure led to more fragmented event segmentation during continuous memory encoding. In contrast, while pupil size and gaze moving speed were sensitive to event boundaries, these metrics were not modulated by prior short video watching, indicating the disruption is specific to the segmentation process itself and not to lower-level boundary detection. Collectively, these findings demonstrate a negative impact of certain short video watching habits on event segmentation and subsequent memory, underscoring the powerful role of platform algorithms in shaping human cognition.

## Introduction

The exponential growth of short video platforms—internationally known as TikTok and its Chinese counterpart, Douyin—has established them as dominant forces in social media, with their user base reaching 1.6 billion in 2024, approximately 21% of whom are adolescents (Retrieved from https://www.businessofapps.com/data/tik-tok-statistics/ on March, 2025).

Short-video platforms used personalized recommendation algorithms to deliver content to target users, enhancing user immersion, potentially leading to short-video addiction^1^, fundamentally altering how people, particularly adolescents, consume information. Given the widespread use of these short-video platforms by populations undergoing critical cognitive development, it is essential to understand their consequences for the foundational mechanisms of learning and memory. This study provides empirical evidence of how this media format influences the ability to segment and encode continuous events, aiming to inform scientifically grounded recommendations for its use.

Previous research has shown that exposure to the fragmented information common on these platforms increase users’ cognitive load^2^ and impair cognitive functions like sustained attention^3^, working memory^2^, prospective memory^4^, time perception^5^, and women’s body satisfaction^6^. We propose that these effects may converge to disrupt a more fundamental memory and learning mechanism: event segmentation. Event segmentation is the fundamental process of parsing continuous experience into discrete, meaningful units^7–9^. This process is crucial for encoding continuous stimuli, such as commercial movies^10–12^, sports footage^13^, and auditory narratives^14^ such as movies or narratives. Critically, effective segmentation during encoding is a strong predictor of subsequent memory recall^15–18^. However, while the broader cognitive impacts of short-video use are emerging, it remains unclear how the platform’s core feature—rapid and frequent context-switching—specifically affects the event segmentation process itself.

Here, we investigate the impact of short-video exposure on event segmentation by testing the central hypothesis that its fragmented format disrupts event segmentation. We focus on two key characteristics of the short-video experience. The first is the fragmented content format, defined by brevity and rapid succession. Videos are typically under 15 seconds, and the user’s ability to switch content at will results in even shorter engagement times. This rapid, user-controlled presentation of abundant and unrelated information establishes a fragmented viewing pattern that contrasts sharply with the sustained cognitive processing required for encoding continuous experiences like watching a feature film or attending a lecture^3^. Such a pattern may increase cognitive load and weaken the encoding of continuous memories^2^.

Because shifts in attention are a primary driver for creating event boundaries during the processing of continuous information^19^, we proposed that the constant, abrupt transitions inherent in short-video viewing disrupt the naturalistic process of event segmentation.

We further hypothesize that this disruptive effect is amplified by the second characteristic: personalized recommendation algorithms. These algorithms predict user interests from Browse history and content tags to curate a continuous stream of tailored videos^20,21^. By tailoring content to users’ preferences, algorithms create highly immersive and pleasurable viewing experiences that reinforce compulsive engagement^22^. Supporting this, neuroimaging evidence indicates that personalized video exposure modulates functional connectivity across large-scale brain networks, suppressing regions involved in cognitive control while activating reward-related pathways ^23,24^, which may underlie the habitual nature of short-video consumption. By hijacking the brain’s reward system, personalization may make it harder for viewers to disengage from the rapid, fragmented flow of content, thereby magnifying the disruptive impact of the constant context-switching on event segmentation. This leads to our second hypothesis: that the tailored, reward-driven nature of personalized content will have a more pronounced disruptive impact on the cognitive mechanisms underlying event segmentation compared to a random, non-personalized sequence of videos.

Previous research on the effects of short video exposure on human behavior has primarily relied on self-report studies^25^. These have indicated that, like reports on smartphone and social media usage, TikTok exposure correlates positively with symptoms of depression and anxiety^26–29^. However, self-reported measures are susceptible to memory biases and social desirability effect. For instance, individuals may experience subjective time distortions when estimating the duration of their online activities^30^, an issue that could similarly affect estimates of time spent on short-video platforms. An alternative research approach involves acutely exposing participants to short videos in a controlled laboratory setting and subsequently observing the carry-over effects on their behaviors. To overcome these limitations and more objectively investigate the cognitive consequences of short-video watching, we employed an integrative research methodology that combined self-reported measures with objective assessments of cognitive functions and memory retrieval.

To test the specificity of this proposed mechanism, we designed two studies contrasting memory encoding for continuous versus discrete information with together of self-reported measures and objective memory retrieval measures. Study 1 used a continuous movie-watching task, which heavily relies on event segmentation for successful memory formation^31–34^. In contrast, Study 2 employed a trial-based static image encoding task, which engages visual memory but does not depend on the same temporal segmentation mechanism^35^. We predicted that if short-video exposure selectively impairs event segmentation, memory performance would be degraded for the continuous movie (Study1) but would remain intact for the discrete images (Study2).

Since memory performance is a relatively indirect measure of the underlying segmentation process, we used eye-tracking, leveraging advanced computational analyses typically applied to neuroimaging data like fMRI^36–41^ and EEG^17^. Leveraging the high temporal resolution and cost-efficiency of eye tracking^42,43^ and building upon Clewett et al.’s findings that pupil dilation reflects changes in event structure elicited by static visual and auditory stimuli^44^, the prior work from our lab that applied this method to identify signatures of event segmentation in continuous memory encoding^45^. We tracked pupil size and eye movement patterns, integrating advanced computational approaches (i.e., *Hidden Markov Models* (HMM)^46^ and *inter-subject correlation analysis* (ISC)^47^) —typically used in neuroimaging—into our eye-tracking data analysis.

More specifically, to capture the potentially subtle disruptions in the dynamic process of event segmentation during continuous viewing, we used advanced eye-tracking analyses, ISC, and HMM, to quantify the dynamic process of event segmentation. We selected Intersubject Correlation (ISC) analysis because it provides a robust measure of shared attentional engagement across viewers during segmentation; lower ISC at narrative event boundaries indicates a divergence in viewing patterns, reflecting disruptions in shared segmentation processes^48^. By applying the HMM—a data-driven event segmentation technique—to eye-tracking data, which includes pupil size and the coordinates of fixation points over time, we determined the model-generated optimal event number, distinctive eye-tracking patterns specific to each event, and the precise timing of event transitions. HMMs are particularly well-suited for identifying latent cognitive states from sequential eye-tracking data, allowing us to quantify the fragmentation of a viewer’s segmentation process.

In summary, we conducted two studies (Study 1, N=113; Study 2, N=60) to assess the potential negative effects of acute short-video exposure on memory encoding in healthy adults (**Figure 1)**. In Study 1 (N=113), we used a four-group, pre-exposure paradigm to test our central hypotheses. Participants’ eye movements were monitored while they encoded a ∼20-minute “*BBC-Sherlock*” episode, a well-established stimulus in event segmentation research, before completing a free recall memory test. Our design was constructed to systematically disentangle the core components of the short-video experience. To isolate the hallmark feature of short video watching—rapid, unpredictable context-switching—we established two short-video conditions. The Short-Random (Short-R) group viewed randomly selected videos, representing our purest test of this disruptive mechanism. In contrast, the Short-Personalized (Short-P) group viewed algorithmically recommended videos from their own accounts, allowing us to examine the potential moderating role of algorithm-driven engagement. These conditions were compared against two long-form video controls that provided critical benchmarks. The Long group watched an unrelated documentary (’BBC-Ocean’), serving as our primary baseline for a standard, continuous viewing experience. Finally, the Schema group watched a related documentary about Sherlock, acting as a positive control to verify that our memory measures were sensitive enough to detect the known facilitatory effects of prior knowledge ^49^. This four-group design allowed us to test for impairments caused by short-video viewing while accounting for the influence of personalization and confirming the task’s sensitivity (**Table 1**). As hypothesized, we found converging behavioral and eye-tracking evidence that short-video exposure disrupts event segmentation during continuous memory encoding. To probe the specificity of this finding, Study 2 (N=60) used a trial-based image encoding task. The absence of a similar deficit in this context suggests that the cognitive impact of short-video viewing may be uniquely disruptive to event segmentation and continuous memory processes.

**Figure 1:**
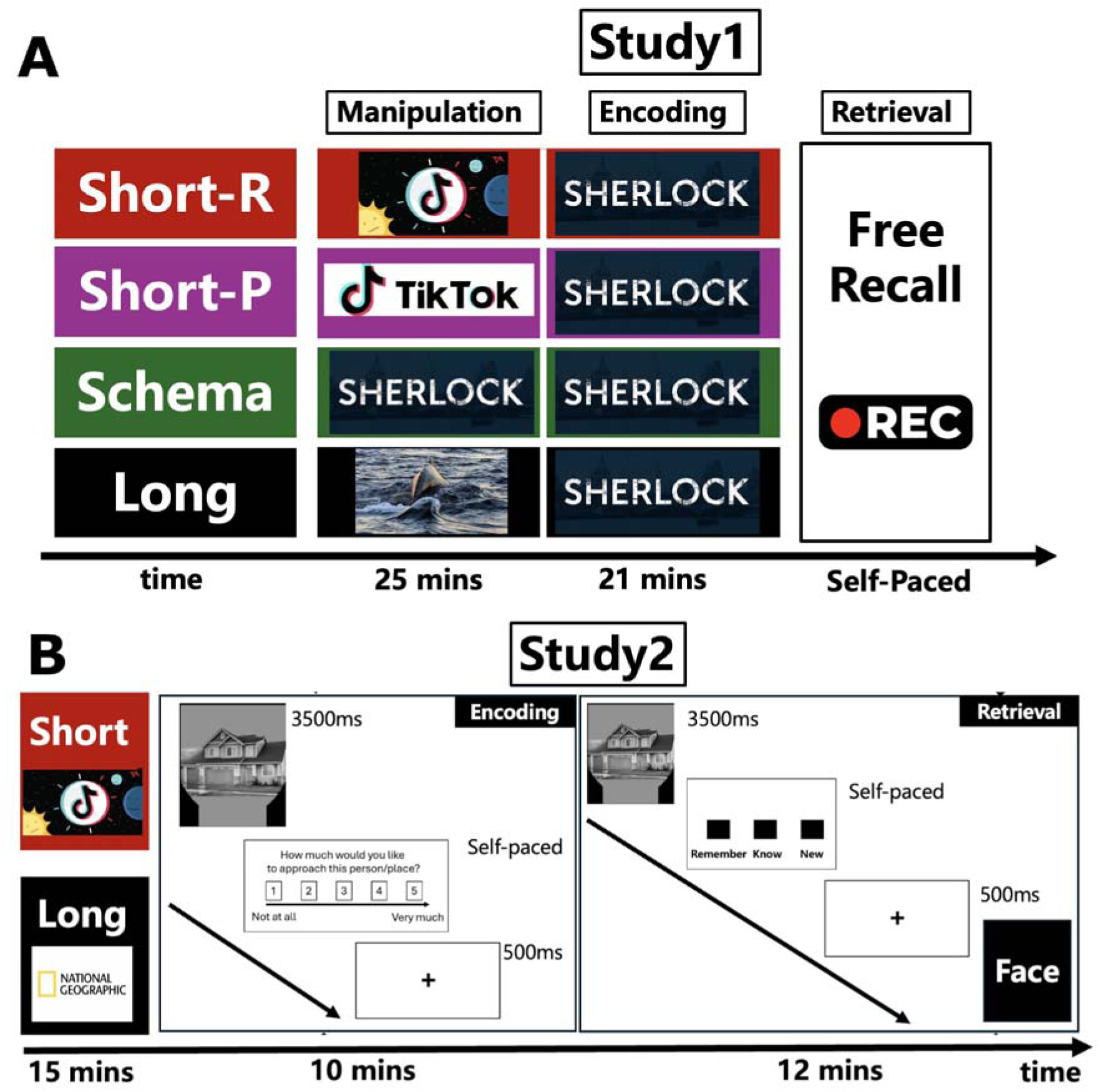
Experimental Design (A) Study 1 Outline. In the manipulation phase, participants were exposed to one of four conditions: randomly selected short videos (Short-R group), personalized short videos (Short-P group), a unrelated segment from the “BBC-Ocean” documentary (Long group), or a related segment from a Sherlock movie (Schema group).

**Table 1.**
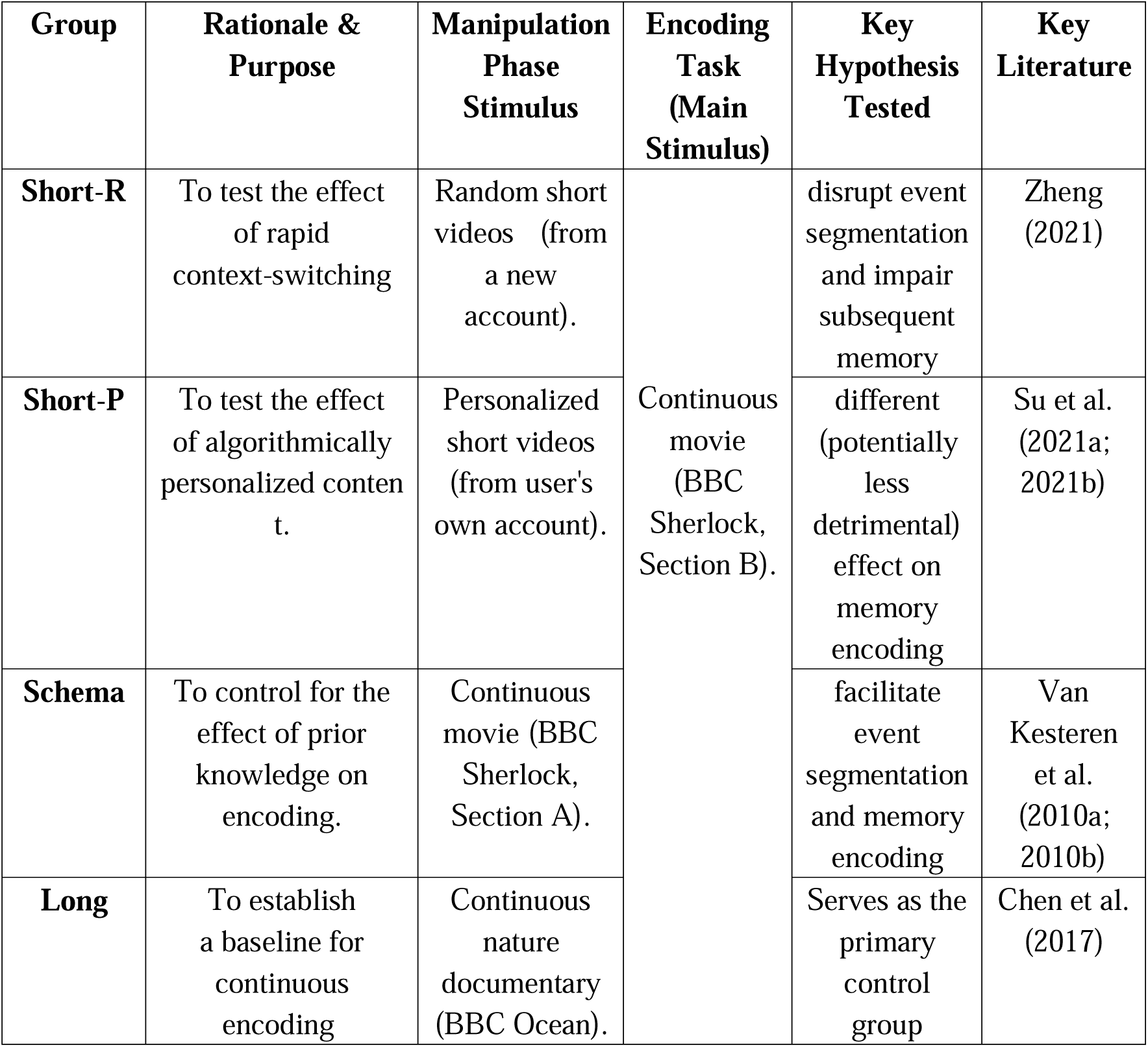
Overview of Experimental Groups, Procedures, and Hypotheses in Study1.

During the encoding phase, all participants viewed the same segment of the Sherlock movie while their eye movements were monitored, and memory was assessed through a subsequent free recall task. **(B) Study 2 Outline.** In the manipulation phase, participants watched either of short videos (Short group) or a documentary from National Geographic (Long group). During the encoding phase, participants rated the approachability of images (faces or houses) in a self-paced task, following the presentation of each image. In the retrieval phase, memory recognition was evaluated using a Remember/Know (RK) paradigm.

## 2. Methods

### 2.1 Participants

#### Participant recruitment and screening

Participants were recruited from the CCNU university community. Eligibility was determined through a pre-study screening process. Prospective participants were required to meet the following criteria: (1) no self-reported history of neurological or psychiatric conditions; (2) a score below 11 on the Beck Depression Inventory-II (BDI-II); (3) self-reported prior experience with using short-video applications (e.g., TikTok); and (4) no prior familiarity with the experimental stimuli (i.e., the’Sherlock’ episode in Study 1 or the specific face set in Study 2). The BDI-II cutoff was established to minimize potential confounds from depressive symptomatology, which can affect memory and attention, and to ensure recruitment from a non-clinical population. Individuals who did not meet all criteria were not invited to the formal testing session. Crucially, all participants who passed this screening and completed the full experimental procedure were included in the final analyses for both studies. No participants were excluded from the final sample after data collection finished.

#### Study1 Naturalistic memory encoding

In Study1, 113 right-handed, healthy undergraduates (73 female, 40 males; average age: 20.42 years, standard deviation: 2.04) were tested. All had normal or corrected vision, no hearing deficits. Recruitment was via advertisements, with compensation ($10 USD) provided for participation in the full session. Each gave written informed consent ahead of the experiments. These were randomly distributed into four groups to receive various experimental manipulations (see the experimental design section for details). The demographics of the groups were: Short-R (28 participants; 11 male, average age = 20.32, SD = 2.08), Short-P (30 participants; 8 male, average age = 20.58, SD = 1.99), Long (28 participants; 9 male, average age = 20.18, SD = 2.11), and Schema (27 participants; 12 male, average age = 20.77, SD = 1.96). No significant differences in age (F=1.145, *p*=0.33) or gender distribution (χ²=2.28, *p*=0.51) were observed between groups. The study protocols were approved by the University Institutional Review Board for Human Subjects.

#### Study2 Picture-based memory encoding

In Study 2, we recruited 60 right-handed, healthy undergraduate students (52 females, 8 males; mean age = 21.7, SD = 1.7), who completed all experimental procedures and were included in the final data analysis. During the manipulation phase, participants were instructed to use their phones to play videos (short or long) within the app. Participants were divided into two age-and gender-matched groups: a Short group (26 females, 4 males; mean age = 21.36, SD = 1.65) and a Long group (26 females, 4 males; mean age = 22.2, SD = 1.71).

Recruitment was via advertisements, with compensation ($10 USD) provided for participation in the full session. The study protocols received approval from the University’s Institutional Review Board for Human Subjects.

#### Sample Size Justification

Given no previous study investigated the effect of short video watching on memory encoding, estimating the precise effect size before collecting data was not possible. The sample size was guided by earlier research on event boundaries that typically engaged 20-30 subjects^10,15,38^, often supplemented by other techniques like EEG and fMRI. A relevant previous study, employing pupil dilation measurements to examine event segmentation, involved 34 participants and posited a need for at least 25 subjects to identify a substantial effect size (d=0.8) under conditions of an α of 0.05 and 80% power^44^. For our Study1, we targeted a minimum of 25 evaluable subjects per experimental condition, ensuring complete datasets of memory and eye movement metrics. All primary and secondary analyses exceeded this participant count, with the minimal sample in the schema group with 27 participants included in the analyses. Study2, we achieved the final sample of 30 participants in our final analyses.

### 2.2 Materials and Design

#### 2.2.1 Study1 Naturalistic memory encoding

##### Experimental design

This Study1 adopted a between-subjects design to investigate the effect of short-video watching on continuous memory encoding. Participants were randomly assigned to one of four experimental groups: (1) the Short-Random (R) group, (2) the Short-Personalized (P) group, (3) the Long group, and (4) the Schema group. The primary independent variable was the type of video content viewed during the manipulation phase. The key dependent variables were (1) memory performance, assessed using a free recall task for a subsequent continuous movie, and (2) eye-movement dynamics during movie encoding, including intersubject correlation (ISC) at event boundaries, latent state patterns derived from a Hidden Markov Model (HMM), and pupil size/gaze moving speed around boundaries.

Manipulation Phase: This phase aimed to influence event segmentation during subsequent video viewing. We randomly divided the participants into four groups: (1) the Short-Random (R) group, (2) the Short-Personalized (P) group, (3) the Long group (watching “Ocean”), and (4) the Schema group (watching the first 25 minutes of “Sherlock”). The viewing duration for the participants in all four groups was 25 minutes. During the manipulation phase, participants in the Short-Personalized group had control over the video playback; they were instructed that they could skip the current video and move to the next one in the downloaded sequence at any time by pressing the designated key. This condition was designed to incorporate an element of user control over the content flow, distinct from the passive viewing in the random condition where videos played consecutively without participant intervention.

Encoding Phase: Immediately following the completion of the manipulation phase, participants proceeded to the memory encoding phase without any delay. During this phase, participants from all groups watched the 21-minute segment from “BBC Sherlock”.

Retrieval Phase (Free Recall): In the free recall phase, participants recounted in detail the content from the encoding phase, emphasizing the depth of recall. The manipulation phase content, while not directly evaluated, was included in assessments to maintain participant focus throughout the experiment. Initial responses were captured via smartphones and later transcribed for detailed memory analysis.

##### Device and Stimuli

All experimental tasks in Study 1 were presented on a 23-inch LCD monitor with a screen resolution of 1280×720 pixels. Stimuli were presented, and behavioral responses were recorded using Experiment Builder software (SR Research Ltd., Ottawa, ON, Canada).

Stimuli for the manipulation phase varied across conditions. Participants in the Short-Random (R) group were exposed to 400 short TikTok clips across themes such as humans, animals, objects, and scenes, each theme contributing 100 clips of 1 to 5 seconds, curated from a neutral account to avoid bias from personalized algorithms. Participants in the Short-Personalized (P) group viewed short videos downloaded from their own TikTok accounts. Participants in the Long group watched a segment of the BBC documentary “Ocean”. Participants in the Schema group watched the first 25 minutes of “Sherlock Season 1 Episode 1 (’A Study in Pink’)”. This episode was previously employed in event segmentation studies^11,12,15^. For the encoding phase, common to all participants, the stimulus was the subsequent 21-minute segment of “Sherlock Season 1 Episode 1 (’A Study in Pink’)”. We selected this episode as it allowed us to adopt previously established event boundaries identified by Chen et al. (2017), ensuring consistency with prior event segmentation research, and because prior studies indicated its capability to evoke distinct, event-specific neural representations ^11,12,15,17^. In total, 46 minutes from the episode were used across the manipulation (schema condition) and encoding phases.

##### Memory measures

Free recall scoring: Participants’ verbal free recall responses were audio recorded for later scoring. The scoring procedure was adapted from Chen et al. (2017), who used the same Sherlock episode stimulus. The original coding scheme segmented the full episode into 50 distinct events, each containing multiple details (approximately 1000 details in total). For the current study, which used a segment corresponding to 20 events from the original study, we created our scoring sheet by extracting these 20 events and their associated details (range: 1-24 details per event) from the Chen et al. (2017) system. This list was translated into Chinese. To enhance scoring accuracy, keywords (e.g., character names, specific actions, dialogue elements) were manually added to the description of each detail by one author (H.X.L.). The original scoring system document from Chen et al. (2017) is available in our OSF repository, along with our adapted and translated version used for scoring.

Rating Process: Two independent raters, blind to experimental conditions, coded the audio-recorded recall protocols. Using the scoring sheet, raters marked each detail as’remembered’ if the participant’s recall accurately mentioned its core content or associated keywords. An event was coded as’remembered’ if one or more of its constituent details were recalled. If no details from an event were recalled, the event was coded as’forgotten’. Inter-rater reliability was assessed on the initial, independent ratings from both coders before any discussion or consensus-building. For the binary measure of whether an event was recalled, agreement was excellent, as indicated by a Cohen’s Kappa of 0.91. For the quantitative measure of the number of details recalled per event, reliability was also high, with an Intraclass Correlation Coefficient (ICC) of 0.86 (calculated using a two-way random effects model for absolute agreement, ICC(2,k)). After this initial reliability assessment, the two raters discussed any scoring discrepancies to achieve 100% consensus. Any scoring disagreements were resolved through discussion to achieve consensus. The final, consensus-based scores were used for all subsequent statistical analyses of the free recall data.

Memory Metrics: Audio recordings from the free recall task were transcribed and assessed by two independent raters. The evaluation of each participant’s recall was based on three metrics: (1) the remember score, reflecting the number of events recalled; (2) the raw detail score, representing the average number of details recalled per event; and (3) the corrected detail score, calculated as the ratio of recalled details to the total details per event, averaged across all events. Additionally, we investigated the serial position effect by calculating the recall probability for each event, considering its position in the original movie’s storyline.

#### 2.2.2 Study2 Picture-based memory encoding

##### Experimental design

In Study 2, we also organized the research into three distinct phases: the manipulation phase, the memory encoding phase, and the retrieval phase (i.e., picture recognition task). Prior to beginning the main experimental tasks, all participants received instructions and completed a brief practice session for the approachability rating task using practice stimuli distinct from those used in the encoding phase.

Manipulation Phase: Following the practice, participants entered the manipulation phase, where they used the Douyin app to watch 15 minutes of either short videos or the long video documentary. For the short video group, participants were instructed to use the TikTok application on their own smartphones as they normally would. This involved naturalistic interaction with the app, including actively scrolling through the feed, selecting videos to watch, and skipping content via user-initiated actions. This approach was chosen to enhance ecological validity, ensuring the exposure involved the typical user control inherent in real-world short-video consumption, before assessing its impact on the subsequent memory task. For both short and long video groups, participants were instructed not to use the comment or like functions of the app and to disable notifications from other apps to avoid interruptions.

Encoding Phase: Following the completion of the manipulation phase, participants proceeded immediately to the memory encoding phase without any delay. Participants underwent an initial encoding session where they were tasked with rating the approachability of 60 faces and 60 non-social images. This activity was structured into two segments separated by a 20-second interval, with each image displayed for 3500 ms followed by a self-paced approachability rating on a scale from 1 (not approachable) to 5 (very approachable). This task masked the true intent of memory encoding, requiring detailed attention to each stimulus. An interstimulus interval of 500 ms was maintained.

Retrieval Phase (Picture Recognition): Memory retention was unexpectedly assessed using the remember/know (RK) paradigm^50^. Participants were presented with the previously encoded images alongside 30 new faces and 30 new non-social images across two blocks, totaling 180 stimuli. They were asked to classify each image as remembered, known, or new.

Each image was shown for 3500 ms, followed by a self-paced response, with trials separated by a 500 ms interval. A brief practice session using unrelated cartoon faces and buildings preceded the testing to clarify the distinction between’remember’ and’know’ responses.

##### Device and Stimuli

Participants watched short/long videos on their own smartphones using their personal Douyin (the Chinese version of TikTok) accounts. This was done to increase the ecological validity of the short-video watching experience. Participants performed a picture-based memory task on a laboratory computer. The memory task was programmed and administered using E-Prime 3.0 (Psychology Software Tools, Pittsburgh, PA). No eye-tracking was performed in Study 2. Stimuli for the manipulation phase consisted of either short videos viewed via the Douyin/TikTok app on participants’ own smartphones or a long video, specifically the documentary “The Enchanting Creatures of the Motuo Forest - Inside the World’s Largest Canyon” from National Geographic. The duration of this phase was 15 minutes. Stimuli for the memory encoding and retrieval phases consisted of a total of 180 images. Ninety images depicted human faces and were sourced from the Chinese Academy of Sciences’ Sinicized Face Emotion Image System. From a pool of 600 grayscale images, we selected 90 faces (45 males, 45 females) displaying positive, neutral, and negative emotions—30 faces per emotion category. Sixty of these faces were employed for memory encoding, and 30 served as distractors during the recognition task. For non-social stimuli, we curated 90 images (30 each of houses, castles, and offices) from pixabay.com and used under Creative Commons licenses permitting reuse for research purposes. These were similarly divided, with 60 used for encoding and 30 for interference, all presented within an oval mask against a black backdrop. Distinct practice stimuli (e.g., cartoon images) were used for the initial practice session of the approachability rating task. Cartoon faces and buildings were used for the practice session before the retrieval phase.

#### Memory measures

Overall recognition memory accuracy was quantified by the net difference between the overall hit rate and false alarm rate [i.e., Overall recognition memory = hit rate (remember + know) − false alarm rate (remember + know)]. Separate metrics for recollection and familiarity were calculated: recollection was defined as the net hit rate for remembered items minus the false alarm rate for those classified as remembered [i.e., hit rate (remember) − false alarm rate (remember)], and familiarity was calculated as the net rate of known responses, adjusted for false alarms [i.e., (hit rate know / (1 - hit rate remember)) - (false alarm rate know / (1 - false alarm rate remember))]. These measures were used to evaluating the impact of experimental variables on long-term memory retention. Given that the images originated from two principal categories—faces and houses—we first computed the Overall Recognition, Recollection, and Familiarity scores separately for each category. This analysis aimed to determine if the act of watching short videos exerted a selective impact on certain categories. Subsequently, we computed all memory metrics without regard to image category. In summary, we derived nine memory metrics for comparative analyses between groups of participants who viewed either short or long videos prior to the encoding phase, and for correlational studies with questionnaire-based assessments of daily short video usage.

### 2.3 Questionnaire measure of short video use

Participants from both Study 1 and Study 2 completed a 20-item Short Video Use Scale at the conclusion of their respective experiments, which was used to assess the degree of participants’ short video usage. This data collection occurred post memory encoding and retrieval phases to prevent participants from inferring the study’s objectives, which could potentially influence their memory performance. Problematic short-video usage was assessed using a scale adapted from the validated Chinese version of the Internet Addiction Test (IAT)^51^. The primary modification involved replacing the term “the Internet” with “Short Videos^52^”. To ensure clarity, the beginning of the questionnaire defined “Short Videos” for participants as encompassing content from platforms such as TikTok (International Version), Douyin (the Chinese version of TikTok), KuaiShou, and the Tencent Short Video Platform. While this adapted scale demonstrated effectiveness in previous research by Su et al. on short-video usage^23,24^, it has not undergone separate, formal psychometric validation. The scale consisted of 20 items rated on a 5-point Likert scale. In our sample, participants typically completed this questionnaire in approximately 2-3 minutes.

### 2.4 Eye-tracking data acquisition

To control for environmental factors, all experimental sessions were conducted under constant artificial illumination within a sound-attenuated, light-controlled room, minimizing potential pupillary changes due to natural light. All participants, regardless of their group allocation, were accommodated in a uniform environment where they were exposed to stimuli on the same model of monitor, with their eye movements being uniformly recorded.

The experimental setting was modified to mitigate any influence from external light by using curtains to block out natural sunlight, ensuring that indoor lighting maintained consistent luminance throughout the day. The precision in monitoring eye movements was achieved through a desk-mounted EyeLink 1000 system (SR Research Ltd., Mississauga, Ontario, Canada), using a 35-mm lens at a high sampling rate of 1000 Hz. A head and chin rest provided by SR Research minimized head movements. Participants were not given specific instructions regarding blinking, allowing for natural viewing behavior during the extended task duration (approximately 45 minutes for the movie encoding). Before the onset of the experiment, participants underwent a briefing on the eye movement calibration and the eye tracker setup, which included adjustments to the camera for optimal pupil visibility and modifications to the infrared sensitivity threshold. Calibration involved directing participants to sequentially focus on the four extremities of the monitor’s display and was initiated by asking them to concentrate on nine randomly ordered black dots across the screen, placed at predetermined locations. The calibration’s extent adapted based on the stimulus task’s screen area, using a 9-point grid. Each point had to be focused upon by the participants until its disappearance, without predicting its trajectory. Post-calibration, the accuracy was verified by reevaluating participants’ ability to fixate on the same nine points, this time assessing the consistency between recorded and actual positions to quantify visual discrepancies.

Participants were required to pass the system’s default calibration validation threshold before beginning the experiment; if the initial calibration was unsuccessful, the procedure was repeated until the criterion was met. Specific quantitative precision and accuracy metrics for each participant’s calibration were not recorded. Eye-tracking data was collected throughout the experiment, including the initial manipulation phase and the subsequent encoding phase (i.e., watching’Sherlock’ movie). Critically, all reported eye-tracking analyses were performed exclusively on the data from the encoding phase. This was because all participants viewed the identical’Sherlock’ movie, providing a common baseline to assess group differences induced by the preceding video manipulation. In contrast, eye-movements during the short-video phase were not directly comparable across participants due to the randomized nature of the video content.

### 2.5 Eye-tracking data preprocessing and analysis

#### Preprocessing

We used Matlab to preprocess the eye-tracking data. Since all videos presented had a fixed, predetermined duration, we segmented the continuous eye-tracking data by extracting the time series corresponding to the exact duration of video presentation during memory encoding. This time-locking procedure yielded data sequences of standardized length for each participant, ensuring the complete temporal information within each segment was retained for subsequent analysis. We also corrected for artefacts due to blinks by linearly interpolating the eye-tracking signal during detected blink periods (typically lasting 100 to 300 ms), culminating in a complete dataset for each participant that included coordinates of screen fixations (X, Y) and pupil diameters.

#### Interject-correlation analyses (ISC) of eye movements

To investigate how event boundaries influence the synchronization of eye movement patterns and the effects of prior exposure to short videos, we calculated the inter-subject correlation (ISC) of eye movements during specific temporal segments around event boundaries (pre-event, during-event, and post-event) for each participant. We selected Intersubject Correlation (ISC) analysis because it provides a robust measure of shared attentional engagement across participants viewing dynamic, continuous stimuli like movies^47,53^. Lower ISC, particularly at narrative transition points (event boundaries), indicates divergence in viewing patterns, potentially reflecting disruptions in shared event segmentation processes. The analysis was conducted by calculating Pearson’s correlation coefficients for vertical gaze positions between each subject and all other viewers, averaging these values to derive an ISC for each individual, and repeating across all participants to establish individual ISC scores. This procedure was also applied to pupil size and horizontal gaze data. A composite ISC was then calculated as the mean of these values: ISC = (ISC_x_ + ISC_y_ + ISC_pupil_) / 3. To assess synchronized eye movements around event boundaries, we calculated Intersubject Correlation (ISC) across three specific time windows relative to each event boundary: Pre-boundary ISC: Calculated using eye-tracking data from the 10 seconds immediately preceding the event boundary (i.e.,-10 seconds to 0 seconds relative to the boundary). Boundary ISC: Calculated using eye-tracking data spanning 5 seconds before and 5 seconds after the event boundary (i.e.,-5 seconds to +5 seconds relative to the boundary).

Post-boundary ISC: Calculated using eye-tracking data from the 10 seconds immediately following the event boundary (i.e., 0 seconds to +10 seconds relative to the boundary).ISC was quantified as the mean correlation of vertical gaze position, horizontal gaze position, and pupil size between pairs of participants within these defined time window. We analyzed ISC variability for each participant across these conditions using repeated-measures ANOVA and conducted between-group comparisons using ANOVA to determine if ISC scores varied among different groups.

#### Eye-tracking-based Hidden Markov Models (HMM)

We adapted Hidden Markov Models, previously used in fMRI^11^ and EEG^17^ studies, to analyze eye-tracking data. To model the underlying structure of event segmentation from the continuous eye-tracking data, we employed Hidden Markov Models (HMMs). HMMs are particularly well-suited for identifying latent cognitive states (i.e., distinct’viewing patterns’ potentially corresponding to perceived events) from sequential data like gaze coordinates^11,38,45^. By analyzing the transitions and properties of these states, HMMs allow us to quantify characteristics such as the fragmentation or coherence of event segmentation during viewing. This method processed three time series: X and Y coordinates of screen fixations and pupil size. Our data-driven event segmentation model could discern specific eye movement patterns and the structural dynamics of events, including the identification of event boundaries. Initially, we explored individual differences in event structures without presetting the number of events (K). We employed the HMM across K values ranging from 15 to 25, allowing the model to select the optimal K based on eye movement data. Previous fMRI studies, using the same movie clip, established a ground truth of 20 events^12^. Consequently, K values below 20 might indicate overlooked responses near annotated boundaries, whereas values above 20 could suggest an over-segmentation of event cognition, with additional boundaries emerging from the eye movement patterns. We further examined the correlation between individual K values and memory performance to affirm the relationship between event segmentation and event memory. As hypothesized, K values exceeding the established ground truth correlated with poorer memory performance. Subsequently, we set the model to a predefined k of 20 to elucidate latent states and the structure of observed events. Using a leave-one-out approach, we initially applied this model with a fixed k to ascertain optimal event boundaries for each participant. We evaluated the model’s fit by comparing the HMM-determined boundaries against those from a null model using within-event similarity metrics. This comparison was statistically analyzed against a distribution generated from 1000 randomized event sequences to assess the significance of the model’s fit. A significant p-value indicated a robust alignment between the observed eye movement patterns and the HMM-derived event boundaries.

#### Pupil size correction for low-level confounds and gaze position

To isolate pupil size changes associated with cognitive engagement, specifically event segmentation, from low-level confounds, we implemented a two-step correction procedure^13^. First, to correct for artifacts related to gaze position, the screen was divided into a grid of spatial bins based on x- and y-coordinates, and the raw pupil area was z-scored within each bin. Second, we employed a multiple linear regression model to partial out variance attributable to low-level sensory features for each participant’s time series. This model included three regressors: (1) Screen Luminance, quantified as the mean grayscale value of each video frame and downsampled to the pupil data’s sampling rate; (2) Motion Energy, calculated as the frame-by-frame absolute difference in luminance to account for visual transients; and (3) Audio Volume, represented by the root-mean-square (RMS) of the audio signal’s amplitude in one-second windows. The residuals from this regression—representing pupil variance after accounting for these sensory confounds—were defined as the “corrected pupil size.” To ensure the robustness of our results, all group-level analyses were performed on both the raw and the corrected pupil size data.

### 2.6 Interpretation of Behavioral and Eye-tracking measures

To clarify the link between our quantitative measures and the underlying cognitive processes, we specified the following interpretive framework. Free Recall (Study 1): The number of recalled events (’remember score’) was interpreted as a direct behavioral index of successful event segmentation. The associated’detail scores’ were used to measure the qualitative richness of the resulting episodic memory traces. Intersubject Correlation (ISC) (Study 1): ISC quantifies shared attentional engagement. We interpret lower ISC at narrative event boundaries as a marker of divergent processing, reflecting a failure of the collective attentional reorientation necessary to update event models. This provides a group-level, eye-tracking-based index of impaired event segmentation. Hidden Markov Models (HMM) (Study 1): The HMM analysis models an individual’s latent event states. We interpret a higher number of inferred states (optimal K) as evidence for more fragmented and unstable internal event models during viewing. This provides an individualized quantification of segmentation coherence, where higher fragmentation indicates disruption. Recognition Memory (Study 2): We used the Remember/Know paradigm to distinguish recollection from familiarity. We interpret recollection (’Remember’ responses) as the retrieval of contextual details, a process reliant on high-quality event segmentation during encoding. We interpret familiarity (’Know’ responses) as a context-free sense of prior exposure, which we predicted would be less sensitive to segmentation quality.

### 2.7 Statistical analyses

All statistical analyses were performed using Python and JASP. To control for multiple comparisons and interactions, we implemented a combination of ANOVA models with post hoc corrections and regression models with interaction terms. Group-level differences in memory performance and eye-tracking metrics were assessed using one-way ANOVAs for each of the primary outcome measures. These models included Group as the between-subjects factor. To control for Type I error due to multiple comparisons, Holm-Bonferroni corrections were applied to the omnibus ANOVA p-values. Significant omnibus results were followed by Tukey’s HSD post hoc comparisons, providing multiplicity-adjusted confidence intervals and effect sizes. Effect sizes were reported as partial eta squared (ηp²) for the ANOVA results, and Hedges’ g or Cohen’s d with 95% confidence intervals for pairwise comparisons. For individual difference analyses, we first calculated within-group correlations. However, to more formally evaluate whether these correlations differed across groups, we conducted an ordinary least squares (OLS) regression model that included independent variable, Group, and their interaction terms. The Short-R group was used as the reference category for the analysis, which allowed for an evaluation of the differential effect of TikTok scores across the other groups. The model’s fit was assessed using F statistics, adjusted R², and regression coefficients with standard errors and confidence intervals. This approach allowed for a more comprehensive analysis of the data, accounting for multiple factors and their interactions within a single framework. The use of mixed ANOVA and regression models addressed the concerns about multiple comparisons and interactions, helping to control for Type I error and ensuring that the complexity of the data was appropriately evaluated.

### 2.8 Data and code availability

Processed behavioral and eye-tracking data have been made available on the OSF platform (https://osf.io/n7k45/?view_only=673d89b23720410799b5daec1f5d511e). Raw data can be accessed by request to the corresponding authors. Due to copyright constraints, original video materials from the study are not available. The analytical scripts and models used in the study are available in the same repository where the data is hosted. Data analyses were performed using MATLAB and Python, with the HMM developed using the BrainIAK toolbox ^54^ and ISC inspired by Perez’s study^14^.

## 3. Results

### 3.1 Random short video watching links daily viewing habits to memory impairment

In Study 1, we first assessed whether self-reported daily short-video usage, as measured by the TikTok score, differed among the four randomly assigned experimental groups. No significant difference was found (F(3,109)=1.63, *p*=0.187; M_Long_=58.43(SD=12.32); M_Schema_=51.26(SD=11.90); M_Short−P_=55.67(SD=10.92); M_Short−R_=52.14(SD=18.13)), indicating that any subsequent group differences in memory were not attributable to pre-existing differences in short video watching habits. We then examined the relationship between TikTok scores and memory performance across all participants. Higher TikTok scores were marginally and negatively correlated with the number of recalled events (r=−0.17, *p*=0.07; **Figure 2A**) but were not significantly correlated with either raw detail recall (r=−0.14, *p*=0.12; **Figure 2B**) or corrected detail scores, which account for variability in available details per event (r=−0.07, *p*=0.41; **Figure 2C**). Group comparisons using ANOVA revealed significant main effects on all three memory metrics: A one-way ANOVA revealed a significant main effect of Group on the number of recalled events (F(3,109) = 5.83, *p* <.001, *p*_Holm_ =.001, ηp² =.138; **Figure 2D**). Significant effects were also observed for raw detail scores (F(3,109) = 3.28, *p* =.024, *p*_Holm_ =.048, ηp² =.083; **Figure 2E**) and corrected detail scores (F(3,109) = 3.00, *p* =.034, *p*_Holm_ =.034, ηp² =.076; **Figure 2F**). The omnibus effect remained significant after Holm correction across the three primary outcomes Tukey-adjusted pairwise comparisons indicated that the Schema group recalled significantly more events than the Short-P group (mean difference = 0.180, 95% CI [0.053, 0.307], *p*_Tukey_ =.002, Hedges’ g = 0.98, 95% CI [0.25, 1.72]), the Long group (mean difference = 0.171, 95% CI [0.042, 0.300], *p*_Tukey_ =.004, g = 0.94, 95% CI [0.19, 1.68]), and the Short-R group (mean difference = 0.148, 95% CI [0.019, 0.277], *p*_Tukey_ =.017, g = 0.81, 95% CI [0.07, 1.55]). For raw detail scores, the Schema group also outperformed the Short-P group (mean difference = 0.71, 95% CI [0.05, 1.36], *p*_Tukey_ =.028, Cohen’s d = 0.75, 95% CI [0.02, 1.48]) and the *p*_Tukey_ group (mean difference = 0.67, 95% CI [0.00, 1.33], *p*_Tukey_ =.049, d = 0.71, 95% CI [−0.03, 1.44]). For corrected detail scores, the Schema group showed significantly higher performance than the Short-P group (mean difference = 0.062, 95% CI [0.001, 0.123], *p*_Tukey_ =.047, Cohen’s d = 0.70, 95% CI [−0.03, 1.42]) and marginally higher than the Short-R group (mean difference = 0.060, 95% CI [−0.002, 0.123], *p*_Tukey_ =.060, d = 0.68, 95% CI [−0.05, 1.42]). No significant differences were observed among the Long, Short-P, and Short-R groups (all *p*_Tukey_ ≥.16, |d| < 0.20).

**Figure 2.**
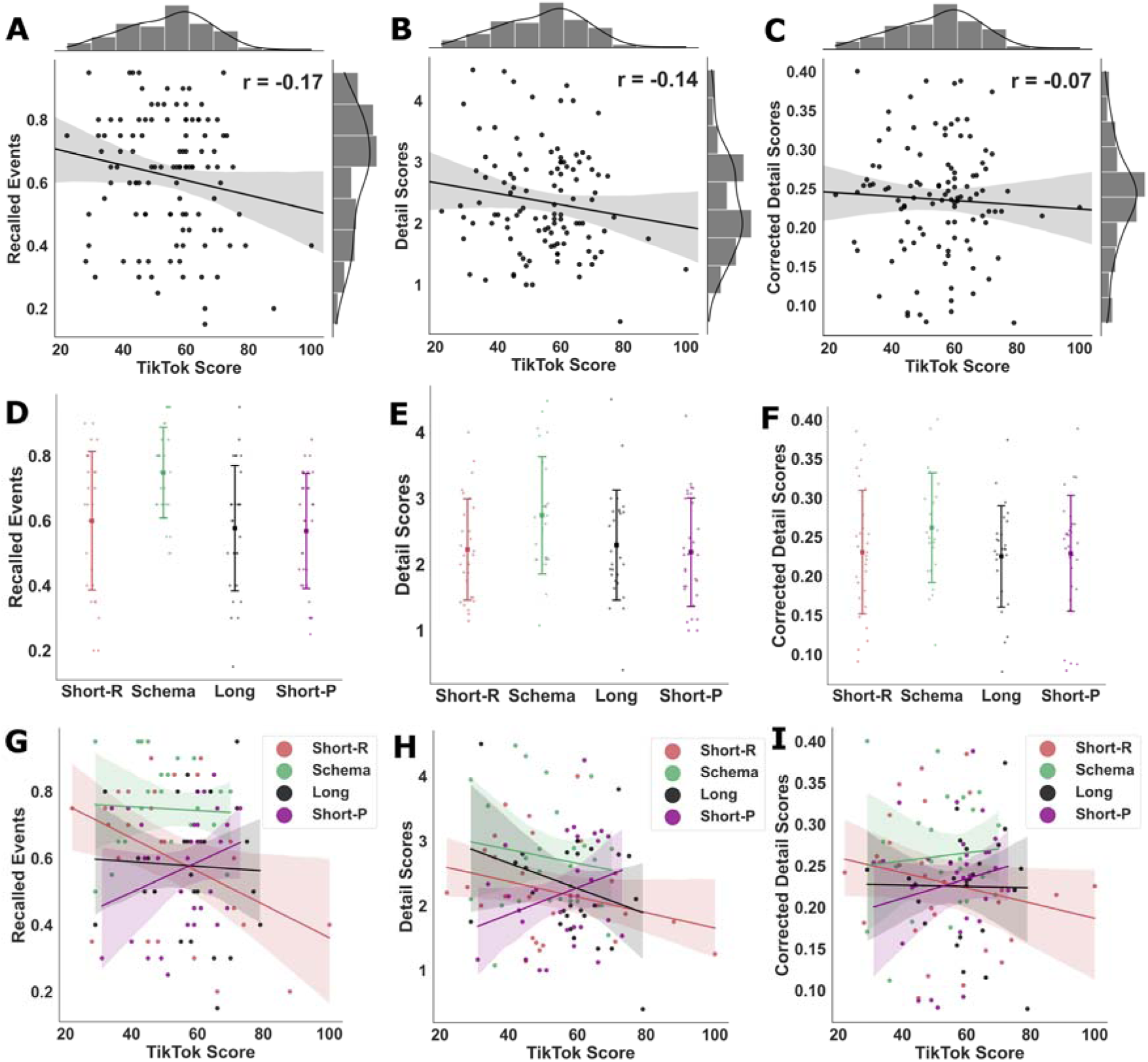
Daily Short Video Viewing Predict Memory Deficits Selectively After Random Short Video Watching. (A-C) Across all participants, higher TikTok scores were marginally correlated with recalling fewer events (r =-0.17, *p* = 0.07) but show no correlation with the detail recall (detail scores: *p* = 0.12; corrected detail scores: *p* = 0.41). **(D-F)** Group-level analysis of memory performance revealed that the Schema group significantly outperformed all other groups in the number of recalled events. Neither the Short-R nor the Short-P groups demonstrated inferior memory performance relative to the Long group across the measures. **(G-I)** Critically, group-specific correlation analyses showed a significant negative relationship between TikTok scores and the number of recalled events exclusively for the Short-R group (r=-0.42, *p*=0.02). This correlation was not significant for the Long, Schema, or Short-P groups.

Finally, we explored correlations between TikTok scores and memory within each group. Critically, we found a significant negative correlation between TikTok scores and the number of recalled events exclusively in the Short-R group (r=−0.42, *p*=0.02; Figure 2G). This relationship was absent in the Long (r=−0.04, *p*=0.81), Schema (r=−0.04, *p*=0.81), and Short-P (r=0.27, *p*=0.13) groups. To test whether the association between TikTok scores and memory performance differed across groups, an OLS regression model with interaction terms was conducted, using the Short-R group as the reference category. The overall model was significant (F(7, 105) = 3.94, *p* <.001), explaining 20.8% of the variance in performance (adjusted R² =.16). A significant main effect of TikTok Score was observed (β = –.005, SE =.002, t(105) = –2.63, *p* =.010), indicating that in the Short-R group, higher TikTok scores was associated with lower performance. In addition, a significant interaction was found between TikTok Score and the Short-P group (β =.010, SE =.004, t(105) = 2.66, *p* =.009), suggesting that the slope of TikTok Score on performance was more positive in the Short-P group compared to the Short-R group. No significant interactions were observed for the Schema or Long groups (all *p* >.20). No other group-specific correlations between TikTok scores and other memory metrics were significant (all *p*s > 0.10; Figure 2H, 2I).

Scatter plots show the line of best fit with a 95% confidence interval.

### 3.2 Disrupted event segmentation after short video watching indexed by reduced inter-subject correlation at boundaries

We employed eye-tracking-based inter-subject correlation (ISC) analysis to investigate the impact of acute short video viewing on subsequent event segmentation during continuous video watching. This ISC calculation incorporated both vertical and horizontal gaze coordinates, alongside pupil sizes, for each participant. ISC was examined under three conditions (**Figure 3A**): 10 seconds preceding the event boundary (pre-boundary condition), 5 seconds before and after the boundary (boundary condition), and 10 seconds after the boundary (post-boundary condition).

**Figure 3.**
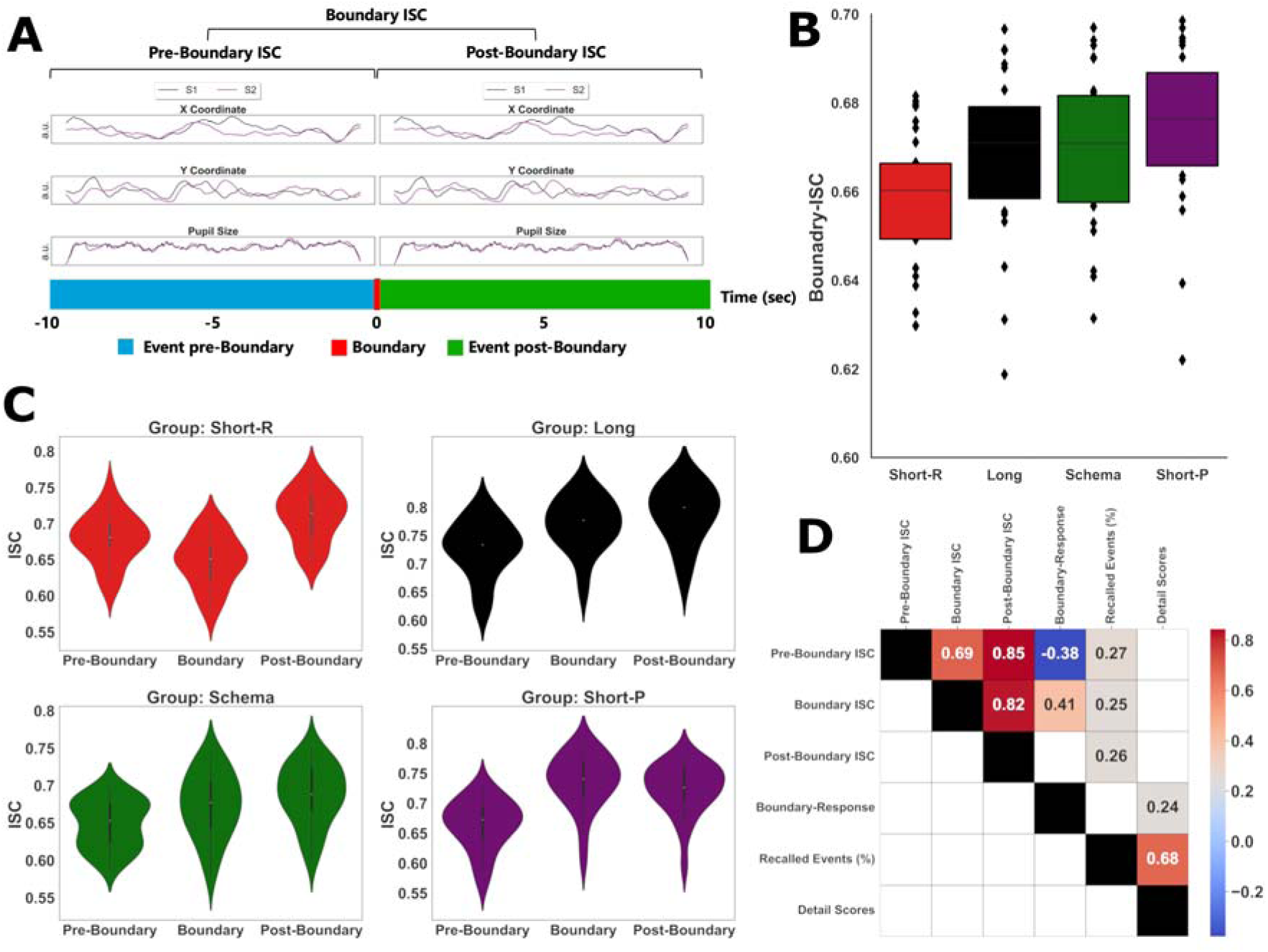
Intersubject Correlation (ISC) Analysis and Event Segmentation During Memory Encoding. **(A)** Displays synchronized eye movements—positions (X and Y coordinates) and pupil sizes—of participants S1 and S2 while they observed two distinct events separated by an event boundary. ISC is quantified as the mean correlation between both vertical and horizontal gaze positions and pupil sizes across participants. Three ISC metrics were analyzed: pre-boundary ISC, calculated during the final 10 seconds before the event boundary; boundary ISC, derived from 5 seconds of eye-tracking data at the boundary; and post-boundary ISC, measured during the initial 10 seconds after the event boundary. **(B)** Participants in the Short-R group, who watched randomly selected short videos, had diminished synchronization of eye movements around event boundaries compared to other groups. **(C)** All groups, except the Short-R group, exhibited higher boundary ISC relative to pre-boundary ISC, a phenomenon defined as the Boundary Response. **(D)** Boundary Response during encoding was predictive of subsequent detailed recall (i.e., detail scores), and that all three ISC metrics were predictive of the ability to recall the number of events (i.e., recalled events (%)).

Group-level differences in boundary ISC were assessed using a one-way ANOVA with Group as the between-subjects factor. The results revealed a significant main effect of Group for the boundary ISC measure (F=4.145, *p*=.008, ηp^2^=0.111; **Figure 3B**), indicating significant differences in performance across groups. Post hoc Tukey tests showed that the Short-P group exhibited significantly higher boundary ISC than the Short-R group (M=0.020, p_Tukey_ = 0.008, Cohen’s d = 0.911, 95% CI [0.145, 1.677]). Other pairwise comparisons did not reach statistical significance (all *p*_Tukey_ > 0.05).

Additionally, within-group dynamic changes from pre-boundary to boundary were analyzed (**Figure 3C**). A mixed repeated-measures ANOVA with time (pre-boundary vs. boundary) as the within-subjects factor and Group as the between-subjects factor showed a significant main effect of time (F = 134.90, *p* <.001, ηp² =.054), indicating that an increase in ISC from pre-boundary to boundary. Importantly, the time × Group interaction was significant (F(3, 100) = 88.06, *p* <.001, ηp² =.106), showing that the extent of increase varied across groups. After Holm–Bonferroni correction for four comparisons, the Long, Schema, and Short-P groups exhibited significant increases in ISC from pre-boundary to boundary (Long: t = 8.29, *p*_Holm_ <.001, Cohen’s d = 1.65; Schema: t = 5.26, *p*_Holm_<.001, Cohen’s d = 1.03; Short-P: t = 13.52, *p*_Holm_ <.001, Cohen’s d = 2.82), whereas the Short-R group showed a significant decrease (t= −8.13, *p*_Holm_ <.001, Cohen’s d = −1.48).

We characterized the increase in ISC from pre-boundary to boundary as a “boundary response” and assessed its association with memory performance. We observed significant positive correlations between the boundary response and the detail score, which measures how accurately participants could recall movie details (r=0.24, *p*=0.01). This finding suggests that the boundary response, absent in the Short-R group, plays a crucial role in linking event segmentation to successful memory encoding. Additionally, the number of events participants recalled (i.e., remember score) demonstrated positive correlations with pre-boundary ISC (r= 0.27, *p*=0.006), boundary ISC (r=0.25, *p*= 0.009), and post-boundary ISC (r = 0.25, *p* = 0.008). We depicted all possible correlations between ISC values and memory performance within the correlation matrix (**Figure 3D**).

### 3.3 Disrupted event segmentation after short video watching revealed by HMM

To objectively characterize event segmentation during naturalistic memory encoding, we applied a Hidden Markov Model (HMM) to participants’ eye-tracking data. We used a leave-one-subject-out cross-validation procedure to determine the optimal number of latent events (K) for each individual. Specifically, for each participant, an HMM was trained on the data from all other participants (N-1) and then used to parse the left-out individual’s time-series data, which comprised pupil diameter and horizontal (x) and vertical (y) gaze coordinates (**Figure 4A**). The optimal K value was identified from a candidate range of 15 to 25. To validate this approach, we constructed a temporal similarity matrix for each participant by correlating their eye-tracking data at every time point with all other time points. Aligning the HMM-identified event boundaries with this matrix confirmed that the model successfully demarcated segments of high within-event pattern similarity, indicating that it captured coherent perceptual events (**Figure 4B**).

**Figure 4.**
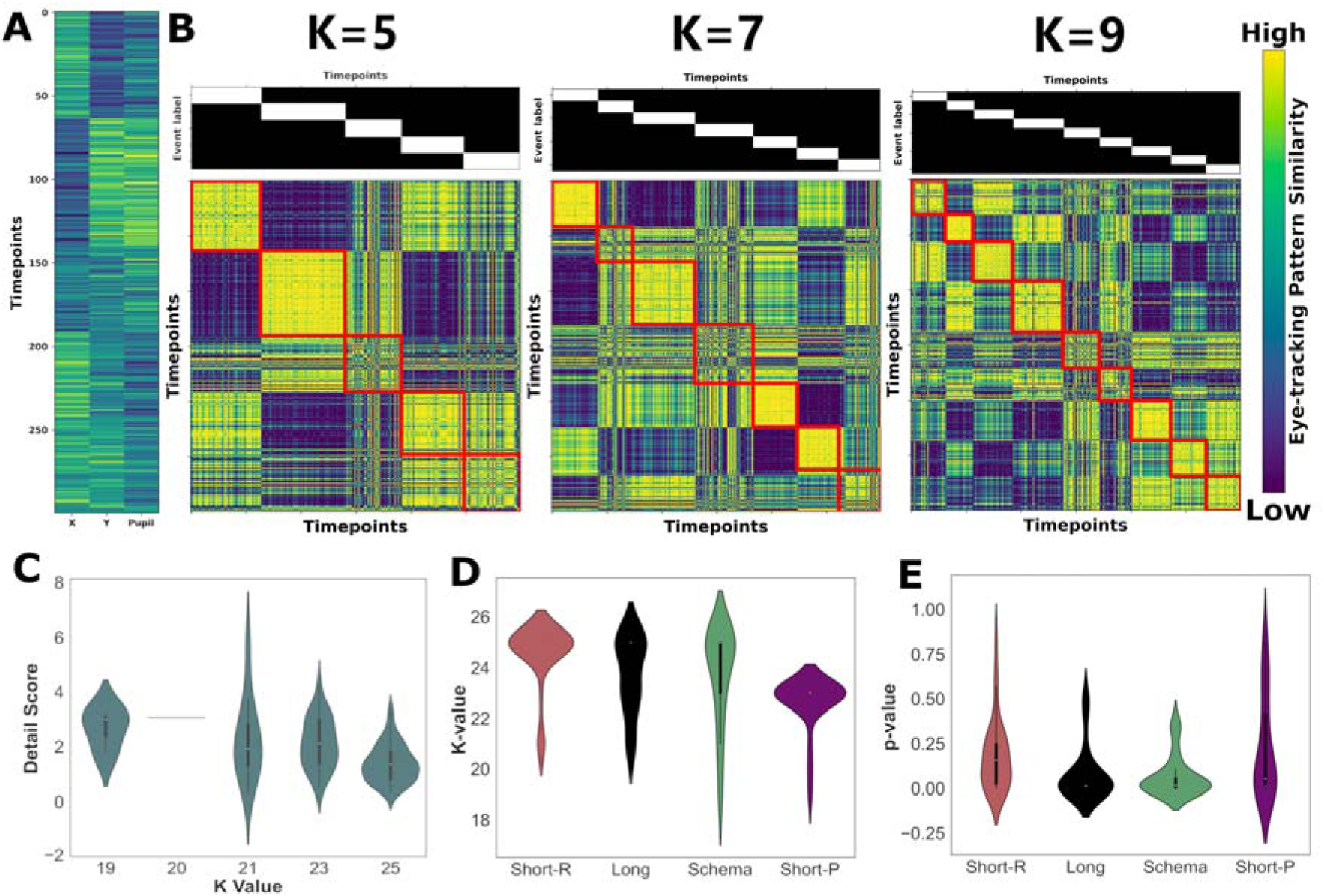
Hidden Markov Model (HMM) analysis of eye-tracking data reveals that disrupted event segmentation is linked to impaired memory recall. **(A)** Schematic of the inputs for the HMM, which include time-series data of pupil diameter and horizontal (x) and vertical (y) gaze coordinates. Data are simulated for illustrative purposes. **(B)** Validation of the HMM approach. HMM-identified event boundaries (red lines) are overlaid on a temporal similarity matrix of eye-tracking pattern similarity. The alignment demonstrates that the model successfully segmented the continuous data into segments of high within-event pattern similarity. The color bar indicates pattern similarity, from low (dark blue) to high (light yellow). **(C)** The optimal number of latent events (K) negatively correlates with memory performance. A higher K, indicating more fragmented event segmentation, is associated with recalling fewer details about the movie. **(D)** Group differences in the optimal number of events (K). The Short-R group exhibited significantly higher K values than the Long group, suggesting more fragmented perception. **(E)** Group differences in HMM model fit, as indexed by p-values. A lower p-value indicates a better fit between the HMM-identified boundaries and the structure of the eye-tracking data. The Schema and Long groups showed a significantly better model fit than both short-video groups.

Having established the validity of this analytical framework, we then proceeded to evaluate the cognitive relevance of the HMM-derived optimal K value. Firstly, we confirmed the cognitive relevance of the HMM-based optimal K value. Across participants, although the optimal K value was not linked to the number of events recalled (i.e., “remember” score; F=0.23, *p*=0.87), it associated with the number of details recalled (“detail” score; F=7.36, *p*<0.001; **Figure 4C**). A linear correlation analysis revealed that the optimal K value negatively predicted the detail score (r=-0.42, *p*<0.001). Subsequent between-group analyses centered on the optimal K value and p-value, assessing the fit between eye-tracking data and model predictions based on HMM-generated event boundaries. A one-way ANOVA revealed a significant main effect of Group on optimal K value (F= 7.26, *p* <.001, ηp² =.179; **Figure 4D**). Tukey-adjusted pairwise comparisons indicated that the Short-P group exhibited significantly lower optimal K values compared to the Short-R group (mean difference = −1.93, 95% CI [−3.00, −0.85], *p* <.001, Cohen’s d = −1.29, 95% CI [−2.08, −0.51]). Other group differences did not reach statistical significance after adjustment. However, comparisons between the Short-P and Long groups (mean difference = −1.11, 95% CI [−0.01, 2.24], *p* =.054, d = 0.75, 95% CI [−0.04, 1.54]) and between the Short-P and Schema groups (mean difference = −1.08, 95% CI [−0.03, 2.20], *p* =.059, d = 0.73, 95% CI [−0.06, 1.51]) showed marginal effects. No significant differences were found among the Long, Schema, and Short-R groups (all *p* ≥.157, |d| ≤ 0.57).

A one-way ANOVA also revealed a significant main effect of Group on p-value (F = 4.14, *p* =.008, ηp² =.111; **Figure 4E**). Tukey-adjusted pairwise comparisons indicated that the Short-P group exhibited significantly higher p-values compared to the Schema group (mean difference = 0.165, 95% CI [0.017, 0.312], *p* =.022, Cohen’s d = 0.84, 95% CI [0.05, 1.62]).

Other pairwise differences did not survive multiplicity correction. Comparisons between the Short-P and Long groups (mean difference = 0.153, 95% CI [0.004, 0.302], *p* =.042, d = 0.78, 95% CI [0.02, 1.57]) approached significance but did not remain reliable after adjustment. No significant differences were found among the Long, Schema, and Short-R groups (all *p* ≥.208, |d| ≤ 0.53).

### 3.5 Unaltered pupil size and moving speed responses to event boundaries across groups

To complement the ISC and HMM analyses, we investigated whether basic eye tracking metrics (i.e., pupil size and gaze moving speed) also reflect event segmentation and were modulated by prior short video watching. First, we analyzed pupil size time-locked to pre-defined event boundaries. Across all experimental conditions, we observed a robust, event boundary-related modulation of pupil size, which peaked at the boundary and subsequently decreased. A repeated-measures ANOVA across five time points (two pre-boundary, the boundary, and two post-boundary) confirmed a significant main effect of time ( F(4,448)=11.69, *p*<0.001, ω^2^=0.095; **Figure 5A**). Post-hoc comparisons revealed a significant decrease in pupil size one second after the boundary relative to the boundary itself (t(112)=6.58, *p*<0.001, Cohen’s d=0.61, 95% CI [0.04,0.09]). We used this post-boundary decrease as a pupillary index of event segmentation. However, a mixed ANOVA testing for a group difference in this index (time points: boundary vs. post-boundary) revealed no significant group-by-time interaction (F(3,109)=0.42, *p*=0.737, ω^2^<0.001; **Figure 5B**), indicating that this pupillary response did not differ across experimental groups.

**Figure 5.**
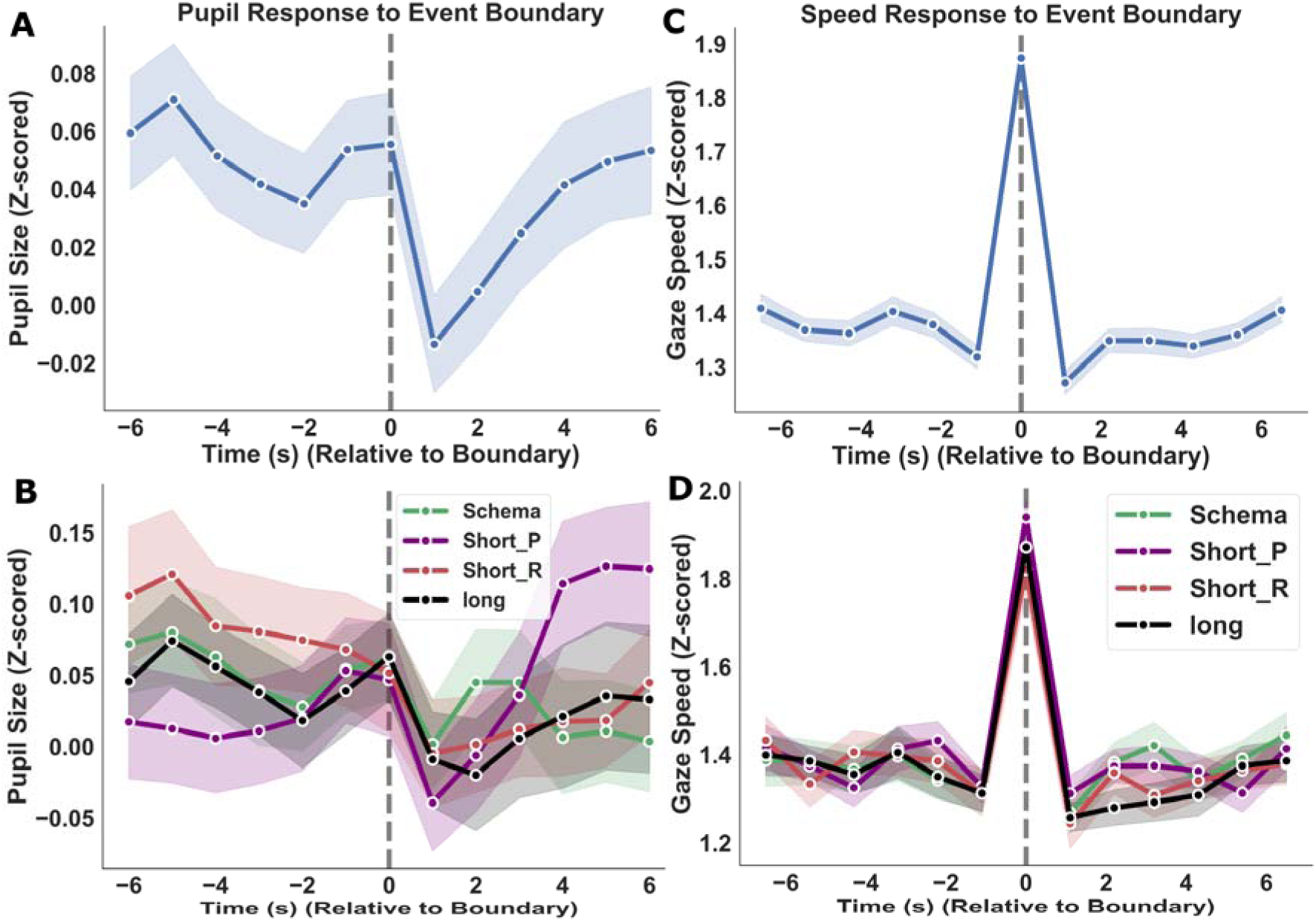
Pupil size and gaze moving speed modulation at event boundaries. **(A)** The time course of mean z-scored pupil size, time-locked to the onset of event boundaries (t=0), averaged across all participants. The plot shows a characteristic peak in pupil size at the boundary, followed by a significant decrease in the subsequent seconds. **(B)** Group comparison of the pupillary responses to event boundaries, No significant differences in this pupillary response were observed across the four experimental groups. **(C)** The time course of mean z-scored gaze speed, time-locked to event boundaries (t=0), averaged across all participants. A prominent and sharp peak in gaze speed is evident precisely at the event boundary. **(D)** Group comparison of the gaze moving speed at event boundaries. The magnitude of the gaze speed increase at event boundaries did not differ significantly across the experimental groups.

A parallel analysis of gaze moving speed across the same five time points also revealed a significant main effect of time, with gaze speed peaking at event boundaries F(4,448)=221.33, *p*<0.001, ω^2^=0.458; **Figure 5C**). Again, the magnitude of this response did not differ between groups, as shown by a non-significant group-by-time interaction in a mixed ANOVA across three key time points (pre-boundary, boundary, post-boundary) (F(6,218)=0.408, *p*=0.873, ω^2^<0.001; **Figure 5D**).

### 3.6 Robustness of eye-tracking results to low-level confounds

To ensure our primary eye-tracking findings were not attributable to low-level stimulus features, we repeated our key analyses after correcting pupil-size data for gaze location, screen luminance, motion energy, and audio volume. A one-way ANOVA revealed a significant main effect of Group on boundary ISC (F= 5.04, *p* =.003, ηp² =.131). Tukey-adjusted pairwise comparisons indicated that the Short-R group exhibited significantly lower boundary ISC compared to the Short-P group (mean difference = −0.025, 95% CI [−0.045, −0.006], *p* =.006, Cohen’s d = −0.93, 95% CI [−1.69, −0.16]). Similarly, the Short-R group showed lower boundary ISC than the Long group (mean difference = −0.021, 95% CI [−0.040, −0.001], *p* =.033, d = −0.75, 95% CI [−1.50, −0.01]) and the Schema group (mean difference = −0.023, 95% CI [−0.042, −0.003], *p* =.014, d = −0.82, 95% CI [−1.56, −0.09]). No significant differences were observed among the Long, Schema, and Short-P groups (all *p* ≥.930, |d| ≤ 0.18).

Similarly, the Hidden Markov Model (HMM) results remained robust. We found a significant main effect of group on the optimal number of hidden states (K-value) (F=3.49, *p*=0.018, ηp²=.095), driven by the Short-R group having significantly higher K values than the Short-P group (mean difference = −2.61, 95% CI [−4.84, −0.38], *p* =.015, Cohen’s d = −0.85, 95% CI [−1.61, −0.09]), which indicates more fragmented event segmentation. No other pairwise comparisons reached significance, including contrasts between the Short-R group and the Long or Schema groups, as well as among the Long, Schema, and Short-P groups (all *p* ≥.124, |d| ≤ 0.83).

### 3.7 No evidence that watching short video disrupts memory encoding of static pictures

We conducted Study 2 with an independent sample of 60 healthy young adults to examine the influence of short video watching on memory encoding and its dependency on the type of stimuli. Our hypothesis was that the picture-based memory encoding task, requiring only brief attention spans and no need for spontaneous event segmentation, would show reduced sensitivity to short video exposure. All participants completed a picture-based encoding-retrieval paradigm using faces and houses as stimuli, along with a self-reported questionnaire assessing daily-life short video usage (i.e., TikTok score). During the manipulation phase, participants were randomly assigned to either watch 15 minutes of short videos or a long video, followed by the encoding phase. The retrieval phase began immediately afterward, and performance was evaluated through separate measures of recognition, familiarity, and recollection for different stimulus types, and collectively for all pictures. No significant differences were found in any of the memory metrics investigated (all *p*>0.10; **Figure 6**, left panels). Furthermore, across all participants, no significant correlations were observed between TikTok scores and memory metrics (all *p*>0.10; **Figure 6**, right panels). Complete statistical results for each group comparison and correlation analysis are available in **Table 2**.

**Figure 6.**
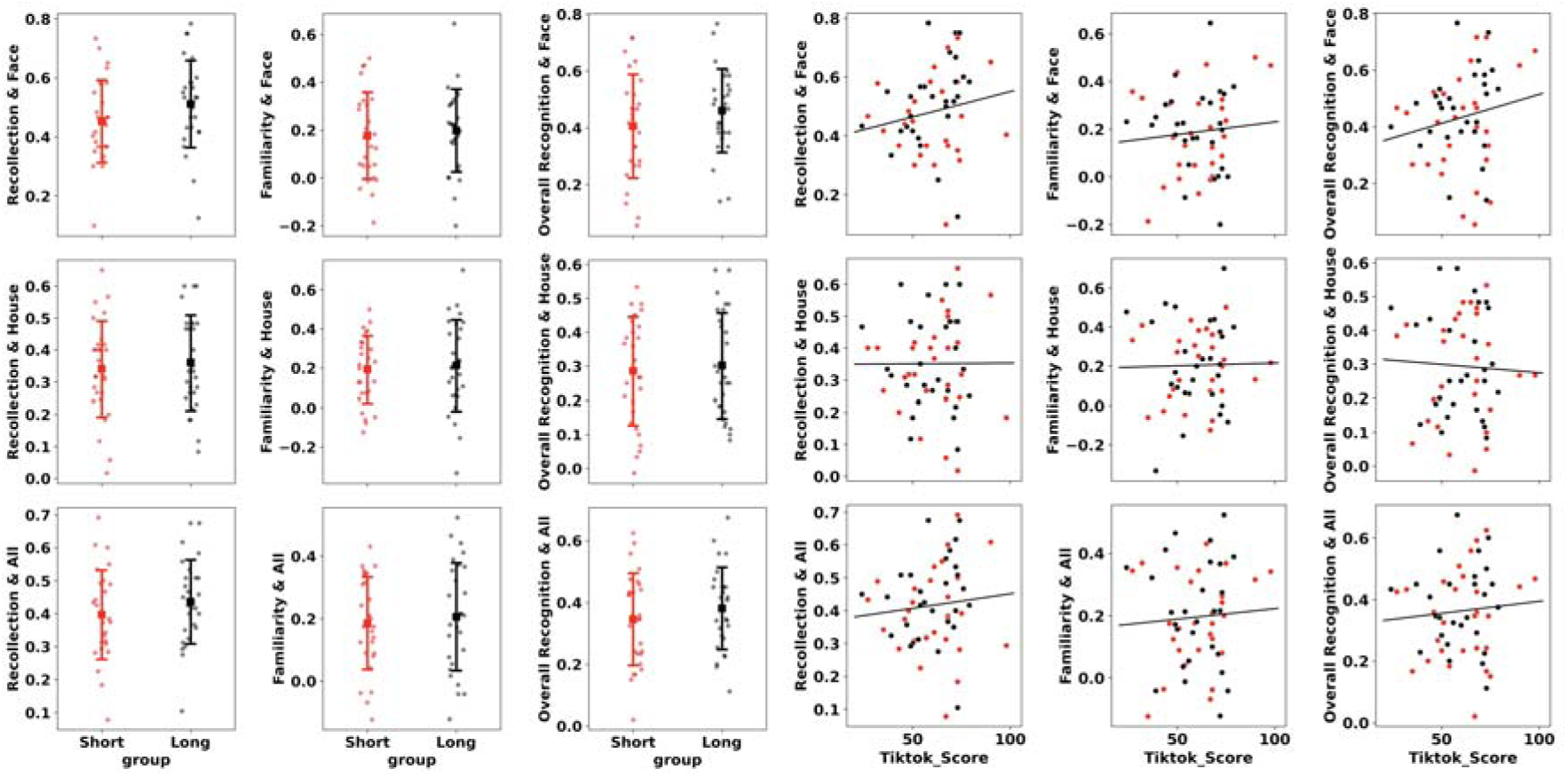
No evidence of impact from acute or daily life short video watching on memory encoding of static pictures. Memory encoding was assessed using a picture-based encoding-retrieval paradigm involving human faces and houses as stimuli. Memory performance was quantitatively assessed through measures of recognition, familiarity, and recollection. The frequency of daily-life short video consumption was quantified via a self-reported questionnaire designed to capture usage intensity (i.e., “TikTok Score”). Left panels: Between-group analysis of memory performance across different stimuli (faces and houses) and memory assessment metrics (recognition, familiarity, and recollection). Right panels: Correlation analysis between TikTok scores and memory performance, encompassing all types of stimuli and memory measures.

**Table 2.** Statistical Analysis of Memory Performance in Study 2.

| Metric<sup>3</sup> | Group |  | T-test<sup>1</sup> |  | Pearson correlation<sup>2</sup> |  |
| --- | --- | --- | --- | --- | --- | --- |
|  | Short | Long | t | p | r | p |
| **Recollection<sub>face</sub>** | 0.45(0.13) | 0.50(0.14) | 1.316 | 0.19 | 0.18 | 0.14 |
| **Recollection<sub>house</sub>** | 0.33(0.14) | 0.35(0.14) | 0.519 | 0.60 | -0.005 | 0.96 |
| **Recollection<sub>all</sub>** | 0.39(0.13) | 0.42(0.12) | 1.013 | 0.31 | 0.10 | 0.44 |
| **Familiarity<sub>face</sub>** | 0.18(0.17) | 0.19(0.18) | 0.242 | 0.80 | 0.07 | 0.54 |
| **Familiarity<sub>house</sub>** | 0.34(0.79) | 0.21(0.22) | -0.818 | 0.41 | 0.14 | 0.25 |
| **Familiarity<sub>all</sub>** | 0.26(0.41) | 0.20(0.16) | -0.680 | 0.49 | 0.15 | 0.22 |
| **OverallRecognition<sub>face</sub>** | 0.40(0.18) | 0.44(0.16) | 0.850 | 0.39 | 0.19 | 0.17 |
| **OverallRecognition<sub>house</sub>** | 0.27(0.16) | 0.30(0.15) | 0.548 | 0.315 | -0.06 | 0.61 |
| **OverallRecognition<sub>all</sub>** | 0.34(0.15) | 0.37(0.13) | 0.814 | 0.419 | 0.07 | 0.58 |

1. Between-group t-tests (t and *p* values) were conducted to explore potential differences in memory metrics between the Short and Long groups. 2. Pearson correlations (r and *p* values) were performed to identify any associations between memory metrics and questionnaire-based measures of daily short video usage (i.e., TikTok score). 3. Overall recognition memory = hit rate (remember + know) – false alarm rate (remember + know). Recollection = hit rate (remember) – false alarm rate (remember). Familiarity = (hit rate know/(1 – hit rate remember)) – (false alarm rate know/(1 – false alarm rate remember)). Subscript indicates that the memory metric was calculated based on pictures of faces, houses, or all pictures used.

## 4. Discussion

Our findings provide robust evidence that watching randomly selected short videos markedly impairs event segmentation during subsequent continuous memory encoding. This cognitive disruption was demonstrated using advanced eye-tracking analyses, including intersubject correlation (ISC) and Hidden Markov Model (HMM) event segmentation model. Specifically, we observed diminished synchronization of eye movements at event boundaries and increased fragmentation in event segmentation following exposure to random, but not personalized, short videos. Importantly, these adverse effects were confined to continuous memory tasks, such as movie watching, and were not observed in discrete, trial-based tasks like picture encoding. These findings indicate that rapid, user-driven content shifts inherent to short-video platforms specifically disrupt event segmentation, thereby altering memory encoding and retrieval processes. This highlights the significant impact of digital media consumption on key aspects of human cognition.

Our behavioral analyses from Study 1 found that acute exposure to short videos, coupled with habitual daily usage, impairs event memory encoding. This suggests that neither self-reported daily use nor acute exposure to short videos independently influences memory encoding. Although previous studies have examined the cognitive impacts of short video watching^5,6,55,56^, none have explored its relationship with human episodic memory encoding. A related study demonstrated that using search engines reduces recall rates of the information itself (i.e., Google effect)^57^. With the rise of short video platforms as a dominant social media form, there is growing interest in their effects on memory. In a parallel Study 2, we found that neither routine daily viewing nor acute exposure to short videos prior to picture encoding affected memory performance. This outcome supports the resilience of picture-based memory encoding to interference from prior short video watching and highlights the advantages of using more ecologically valid stimuli and paradigm in studies of human cognition^58,59^, specificity pointing to the disruption of event segmentation as the core underlying mechanism. Our results underscore the necessity of using ecologically valid paradigms and investigating the interaction between acute exposure and habitual short video watching to fully characterize its cognitive consequences.

Our eye-tracking analysis, using advanced HMM and ISC methods, provide direct evidence that short-video exposure disrupts the mechanics of event segmentation. We propose this disrupt stems from the watching pattern of short videos fragmenting sustained attention. Participants who viewed short videos prior to memory encoding demonstrated disrupted event segmentation, as evidenced by abnormally high numbers of segmented events and decreased synchronization of eye movements near event boundaries. Specifically, after participants watched randomly selected short - videos, during the memory - encoding process, the synchrony of eye movements at event boundaries weakened, and the fragmentation of event segmentation increased. Crucially, these eye-tracking metrics were predictive of subsequent memory performance; greater segmentation fragmentation and lower synchrony were associated with poorer recall. Both neural markers of event segmentation, derived from HMM and ISC, have demonstrated comparable cognitive relevance^10,15,48^ and have been used to investigate developmental changes^38,60^, anticipation^36^, schema^61^ in event segmentation. Our study contributes a novel dimension to Event Segmentation Theory (EST)^7–9^ by applying it to understand the carry-over effects of a prevalent real-world behavior—short-video watching—on subsequent cognitive processing. According to the EST, individuals spontaneously segment their ongoing experiences into discrete events, resulting in event boundaries, which are the time points at which one event ends, and the memory encoding begins^62^. This is a process crucial for the long - term storage of event representations. Event segmentation gives priority to information located at the boundaries, manifested as better memory performance for information at these boundaries. A study found that after participants watched a movie and took a memory test, information at event boundaries was more easily recognized than that at non - event boundaries^63^. Additionally, people are more likely to integrate information within an event boundary into a whole. Compared with recall within the same event, cued and sequential recall across event boundaries is lower^64,65^.

Prediction error plays a crucial role in event segmentation. Our brains constantly predict what will happen next^66^. When the prediction does not match the reality, a prediction error occurs^67^. While EST traditionally focuses on event boundaries triggered by the immediate perceptual stream and internal prediction errors^68,69^, our work demonstrates that segmentation is also susceptible to the residual cognitive states induced by prior cognitive activities (i.e., short video watching in this study). We propose that the frequent and rapid context shifts inherent to short-video watching degrade sustained attention^3,70^. This fragmented attentional state appears to carry over to subsequent tasks, impairing the temporal integration necessary for robust event segmentation. This interpretation aligns with established links between attentional control^71^ and the ability to segment ongoing events and form coherent episodic memories.

Our comprehensive eye-tracking analyses, integrating advanced models (i.e., ISC and HMM) with basic metrics (i.e., pupil size and moving speed), suggest the cognitive mechanisms underlying the disruption in event segmentation following short-video watching. The results indicate that this disruption originates at a high-level conceptual stage rather than from alterations in low-level perceptual processing or arousal. Consistent with prior work showing that HMM-based event segmentation models are robust to low-level visual features^13,45^. Specifically, our HMM analysis identified a greater number of latent eye-movement states post-exposure. This suggests that participants formed more fragmented, internally generated event representations. In contrast, we observed no significant changes in pupil size or moving speed surrounding predefined event boundaries, indicating that arousal and basic perceptual responses to major event transitions remained stable. This dissociation—an increase in latent event boundaries without a change in physiological responses at explicit boundaries—provides novel experimental evidence for a specific mechanism of altered event segmentation. Whereas other manipulations might shift boundary perception (e.g., anticipation^36^) or alter event duration (e.g., in developmental stages^18,38^), short-video exposure appears to uniquely promote the fragmentation of continuous experience. This interpretation of fragmented segmentation supports our central argument of disrupted sustained attention and aligns with broader findings on the cognitive consequences of short video watching.

The divergent results of Study 1 and Study 2 provide compelling evidence that short-video exposure does not cause a global memory impairment, but rather selectively disrupts the process of event segmentation. The continuous movie task in Study 1 required participants to actively segment a temporal stream of information, a process our results show is vulnerable to disruption following exposure to fragmented content. In contrast, the discrete image-encoding task in Study 2 relied more on item-specific visual processing, which does not depend on temporal segmentation. The null result in Study 2 thus serves as a crucial control, indicating that the impairment observed in Study 1 is tied specifically to the demands of processing continuous, unfolding events. We propose this stems from the viewing pattern fragmenting sustained attention and imposing cognitive load, thereby disrupting the event segmentation crucial for encoding temporally unfolding information. In contrast, Study 2 found no effect on memory for static images, likely because encoding discrete items relies more on visual perception and selective attention and less on the temporal integration provided by event segmentation. Therefore, the detrimental impact of such short-video viewing appears specific to cognitive processes dependent on segmenting continuous experience, highlighting the vulnerability of attentional continuity and temporal integration mechanisms to short-video watching involving rapid context shifts.

Our findings reveal a crucial distinction: the observed impairments in memory encoding and event segmentation were driven specifically by exposure to randomly selected, not personalized, short videos. This result establishes the content generation algorithm as a key boundary condition for the cognitive impact of short-video watching. While previous research has predominantly focused on the addictive potential of personalized algorithms^23,24^, our study provides robust behavioral and eye-tracking evidence that non-engaging, random short videos can be uniquely detrimental to continuous memory processes and event segmentation. A plausible mechanism is that random videos, often misaligned with user interests, may decrease attentional engagement. Reduced attention is known to disrupt event segmentation^55^, which in turn impairs memory encoding. Our findings have significant implications for educational contexts. Effective learning from a lecture or a textbook depends on the ability to segment a continuous flow of information into a coherent structure of concepts. The segmentation impairment we observed suggests that habitual exposure to short-form media may hinder a student’s ability to build these structured mental representations from temporally extended educational materials, potentially impacting comprehension and long-term retention. Although our study focused on university students, these findings raise concerns regarding potential relevance to other age groups heavily engaged with short-form video, such as adolescents, and may extend to other cognitive tasks requiring sustained attention and temporal integration, such as complex problem-solving and in-depth reading comprehension.

Our study has several limitations that warrant acknowledgment. Firstly, no eye-tracking data were collected when participants engaged in the picture-based memory encoding task (i.e., Study 2). However, this was due to the methods used for analyzing eye movements being suitable only for capturing cognitive processes during naturalistic memory encoding tasks.

Consequently, it was not possible to compare eye-tracking metrics between Studies 1 and 2. Secondly, we recruited participants who are typical daily users of short videos; specifically, we did not recruit those with a severe addiction to short videos. Future studies could include a cohort of heavy users in similar study design. Thirdly, our study did not include direct assessments of individual cognitive abilities, such as attentional control and working memory capacity. While we assumed that random assignment to experimental groups would mitigate systematic group differences, an important avenue for future research will be to explore these individual differences. For instance, individuals with higher baseline working memory capacity may be more resilient to the fragmenting effects of short-video exposure. Fourthly, the use of foreign film material (Sherlock) with Chinese participants introduces a potential issue of cultural familiarity. However, the between-subjects design, where all groups viewed the same encoding material, helps to minimize this factor’s impact on our main conclusions. Future research should still explore how short-video exposure might differently affect participants depending on their familiarity with the cultural context of the stimuli. Fifthly, a potential limitation of our study is the inherent difference in content between the short-video and long-video conditions, such as topic and format. One might argue for a more controlled comparison, for instance, by segmenting the documentary used in the long-video condition into shorter clips. However, we contend that such a design, while controlling for content, would fail to capture the ecological validity of modern short-video platforms. The core experience of platforms like TikTok or Douyin is the rapid consumption of unpredictable, thematically diverse, and context-free content. Our primary goal was to investigate the cognitive impact of this specific, real-world viewing pattern. Therefore, we intentionally prioritized ecological validity in our design, accepting that content variability is an intrinsic feature of the phenomenon, rather than a confound. Finally, we used commercially available movies as stimuli in Study 1, which may have masked memory impairments in participants who viewed short videos prior to memory encoding, due to the high attractiveness and emotional content of the movies. Future research could employ more emotionally neutral and educationally relevant video clips, such as recordings from real-life educational lectures^48,72^, as memory encoding material.

In conclusion, our findings demonstrate that watching randomly selected short videos significantly impairs event segmentation during continuous memory encoding. Importantly, these effects are absent in discrete memory tasks, underscoring the unique challenges posed by processing continuous information. Given that self-report studies have predominated in short video research^25^, we advocate for interdisciplinary efforts, particularly involving cognitive psychologists and neuroscientists, to further investigate and elucidate the changes in event segmentation caused by short video watching across varied demographics, with a special focus on children and adolescents.

## Competing Interests

The authors declare that they have no known competing financial interests or personal relationships that could have appeared to influence the work reported in this paper.

## Acknowledgments and Funding

W.L was supported by the National Natural Science Foundation of China (grant No. 32300879 and No. W2421004), Humanities and Social Sciences Fund, Ministry of Education (grant No. 22YJCZH109). X.H was supported by the Knowledge Innovation Program of Wuhan-Shuguang Project (2023020201020384).

## Contributions

W.L. and X.H conceived the Study; J.S.L and H.X.L collected the data; W.L. and J.S.L analyzed the data; W.L. and H.X.L prepared the first draft. W.L. and X.H reviewed and edited the manuscript, provided supervision, and obtained funding.

## Notes

### Competing Interest Statement

The authors have declared no competing interest.

### Summary of Updates

The primary revisions that have enhanced the manuscript include: 1. Enhanced Methodological Rigor via New Control Analyses: We have strengthened our pupillometry analyses by incorporating robust controls for low-level visual features and gaze location, as astutely suggested by the reviewers. This new analysis confirms that our findings are not driven by simple changes in arousal or low-level visual features, but rather by more nuanced disruptions in cognitive processing. 2. Strengthened Theoretical Framework and Interpretation: The Introduction and Discussion sections have been substantially revised to better contextualize our study within Event Segmentation Theory (EST). We now offer a more nuanced interpretation of our findings, clarifying how short-video watching can externally modulate the event segmentation process, a novel contribution to the field. 3. Improved Clarity and Justification of Experimental Design: We have thoroughly edited the manuscript to clarify key terminology and improve the overall flow of the arguments. A new table has been added to explicitly link each experimental condition to its theoretical rationale and specific hypothesis, and we have provided a clearer justification for the inclusion of both continuous and discrete memory tasks.

